# InstantDL - An easy-to-use deep learning pipeline for image segmentation and classification

**DOI:** 10.1101/2020.06.22.164103

**Authors:** Dominik Waibel, Sayedali Shetab Boushehri, Carsten Marr

## Abstract

**Motivation:** Deep learning contributes to uncovering and understanding molecular and cellular processes with highly performant image computing algorithms. Convolutional neural networks have become the state-of-the-art tool to provide accurate, consistent and fast data processing. However, published algorithms mostly solve only one specific problem and they often require expert skills and a considerable computer science and machine learning background for application.

**Results:** We have thus developed a deep learning pipeline called InstantDL for four common image processing tasks: semantic segmentation, instance segmentation, pixel-wise regression and classification. InstantDL enables experts and non-experts to apply state-of-the-art deep learning algorithms to biomedical image data with minimal effort. To make the pipeline robust, we have automated and standardized workflows and extensively tested it in different scenarios. Moreover, it allows to assess the uncertainty of predictions. We have benchmarked InstantDL on seven publicly available datasets achieving competitive performance without any parameter tuning. For customization of the pipeline to specific tasks, all code is easily accessible.

**Availability and Implementation:** InstantDL is available under the terms of MIT licence. It can be found on GitHub: https://github.com/marrlab/InstantDL

## 1 Introduction

Deep learning has revolutionised image processing (LeCun, Bengio, and Hinton 2015). On specific tasks such as cell segmentation (Caicedo, Roth, et al. 2019; Hollandi et al. 2020), cell classification (Cireşan et al. 2013; Buggenthin et al. 2017; Christian Matek et al. 2019) or in-silico staining (Ounkomol et al. 2018; Christiansen et al. 2018), deep learning algorithms have led to breakthroughs in biomedical image analysis. They now achieve higher accuracy than trained experts (Esteva et al. 2017; McKinney et al. 2020; Christian Matek et al. 2019) and outperform humans at data processing speed and prediction consistency (Tschandl et al. 2019; Liu et al. 2019).

However, machine learning algorithms are mostly developed to solve one specific problem. Moreover, applying them often requires a strong computer science and machine learning background. This makes it difficult for biomedical researchers to identify and use appropriate algorithms. Standardizing and generalizing deep learning methods and thereby reducing the necessary expertise in computer and data science will lead to broader application and acceleration of research.

We thus provide InstantDL, an easy-to-use deep learning pipeline, which automates pre- and post-processing for convenient application. It can be used for semantic segmentation (i.e. the classification of each pixel into a particular class), instance segmentation (i.e. the detection and classification of objects), pixel-wise regression (i.e. in-silico staining) and image classification (i.e. to discriminate cancerous from healthy cells). Only eleven parameters have to be set in order to run the pipeline. Since biomedical datasets are often sparsely annotated, as manual annotation is laborious and costly, we provide pre-trained models for efficient transfer learning.

InstantDL is benchmarked on seven publicly available (see section 6) datasets: multi-organ nuclei segmentation, nuclei detection in divergent images, lung segmentation from CT scans, in-silico prediction of a mitochondrial and nuclear envelope staining, cell classification in digitized blood smears, and cancer classification on histopathology slides. Without any hyperparameter tuning, we achieve competitive results.

InstantDL is aimed at users with a basic understanding of machine learning, knowing suitable loss-functions and understanding how to split data into train and test set. The code however is open source and well documented for those who want to customize the pipeline to their needs. By providing a debugged, tested, and benchmarked pipeline we help reduce errors during code development and adaptation, and contribute to reproducible application of deep learning methods.

## 2 Methods

InstantDL offers the four most common tasks in medical image processing: Semantic segmentation, instance segmentation, pixel-wise regression, and classification (Litjens et al. 2017; Maier et al. 2019). In the following we describe the algorithms implemented in InstantDL to address these tasks and how they can be applied within the pipeline.

### Semantic segmentation

One of the standard approaches for detecting image patterns is semantic segmentation (Minaee et al. 2020). For each pixel in the input image a class label is predicted by the algorithm. The U-Net (Ronneberger, Fischer, and Brox 2015) is a commonly used architecture for semantic segmentation with numerous applications in biomedicine (Falk et al. 2019; Isensee et al. 2018). It consists of a symmetric contractive path to capture context and an expansive path to capture fine localizations (Ronneberger, Fischer, and Brox 2015). The U-Net outputs continuous values between 0 and 1 for each pixel, which can be interpreted as probabilities to belong to a given class. InstantDL allows for two classes (background vs. foreground) and thresholds the output using Otsu’ method (Otsu 1979; Caicedo et al. 2017). In InstantDL we made minor changes to the architecture from (Ronneberger, Fischer, and Brox 2015): (i) we use padded convolutions to receive the same output and input dimensions (ii) and have implemented dropout layers in the encoder to enable uncertainty estimation.

### Instance segmentation

Instance segmentation is used to detect objects (instances) within an image (Minaee et al. 2020; Caicedo et al. 2017). We implemented the Mask-RCNN (Kaiming et al. 2017) in InstantDL for this task. It first detects a bounding box for each object in the image and then performs a segmentation in each bounding box. Our Mask-RCNN is based on a ResNet50 from Abdullah’s (2017) implementation. The Mask-RCNN requires instance level ground truth: For each image in the training set, a set of labels has to be created, each containing a segmentation mask of one single instance. An algorithm to create instance level ground truth from binary semantic segmentation ground truth is provided as a jupyter-notebook with the pipeline.

### Pixel-wise regression

Tasks where no pixel-wise class labels but a continuous pixel value is desired (such as in-silico staining, (Ounkomol et al. 2018)) are called pixel-wise regression. InstantDL uses the same U-Net implementation as for semantic segmentation. The only difference is that the U-Net output is not interpreted as probabilities to belong to one or another class, but is regarded as a regression. We thus use continuous labels for training and the mean-squared-error as regressive loss as proposed previously (Ounkomol et al. 2018).

### Image classification

Here, the task is to classify each image into one of a specific number of given classes. For this task a residual network (He et al. 2016) is implemented. These architectures are widely used for biomedical image classification (Brinker et al. 2018; Liu et al. 2019). Residual networks use residual blocks, a reformulation of layers as learning residual functions, which enable the use of many layers, while ensuring convergence (He et al. 2016). We use a slightly modified ResNet50 with 50 layers (*Keras-Contrib* n.d.) in InstantDL, where we have added dropout layers to enable uncertainty estimation.

## 3 The InstantDL pipeline

### Data preparation

Data has to be manually split in a train and a test set (see Fig. 1A) according to the user’s hypothesis: For one dataset a random split might be suitable (Waibel et al. 2018), while for others a split on patient or tissue slide level is methodically appropriate (Christian Matek et al. 2019). InstantDL can process stacked images, enabling the prediction from multiple channels. Input and ground truth images must have the same filename including the file ending.

**Figure 1.**
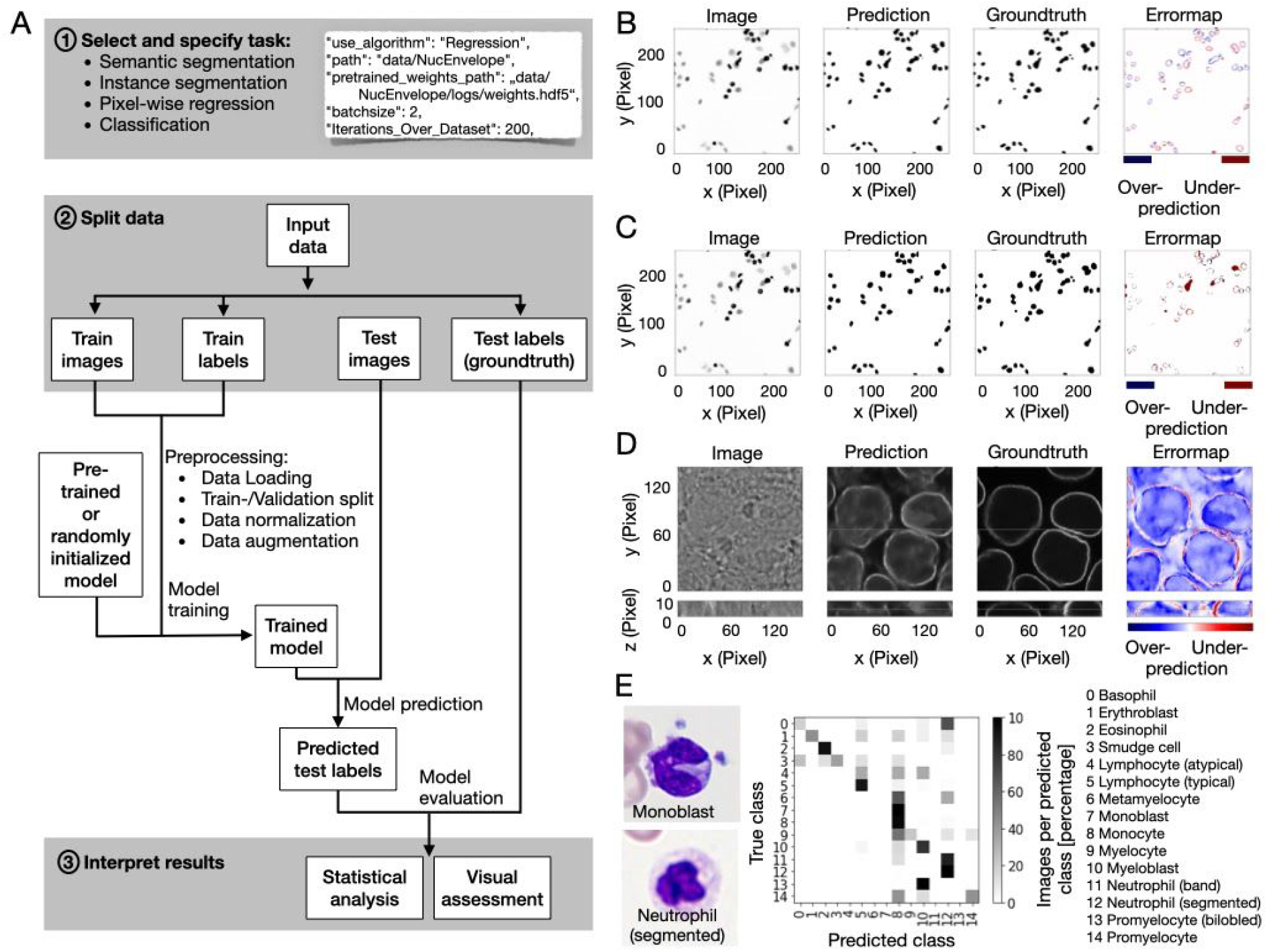
InstantDL provides an easy to use framework for the analysis of biomedical images. (A) Flow diagram of the pipeline with indicated user action highlighted in gray. (1) One out of four tasks (semantic segmentation, instance segmentation, pixel-wise regression, classification) is selected by the user. Up to ten parameters can be set in the configuration file to adapt the pipeline to the task. A code snippet illustrates task selection and six of the ten parameter settings in the configuration file: selected task (“use_algorithm”), path to folder (“path”), if pretrained weights should be used the path to these (“pretrained_weights_path”) should be set, batch size (“batchsize”) and epochs chosen (“Iterations_Over_Dataset”). (2) Input data is split into train and test sets. The user specifies these by putting the data in the corresponding folders. After executing the python configuration file the pipeline will automatically load the data from the train folder, create a 20 percent validation split, normalize and augment the data (see Methods for details). Training is initiated with either a pre-trained or randomly initialized model. After training, the model predicts test labels: segmentation masks, pixel values or labels for the images in the test set according to the chosen task. (3) Results can be interpreted by the user via statistical and visual assessment of the predicted outcome by comparing it to the ground truth in the test set. (B) Example output for a 2D semantic segmentation task: Cell nuclei in a brightfield image (left) are segmented with InstantDL (Prediction) using the U-Net, and compared to the original annotation (Groundtruth). The Errormap indicates over- and under-predicted pixels. The image is part of the 2018 Kaggle nuclei segmentation challenge dataset (Caicedo, Goodman, et al. 2019). (C) Example output for a 2D instance segmentation task (same image as in (B)): A binary mask is predicted for each object in the image using InstantDLs Mask-RCNN algorithm and compared to the groundtruth. (D) Example output for a 3D pixel-wise regression task using a U-Net. From stacks of bright-field images (Image) (Ounkomol et al. 2018) the pipeline predicts a nuclear envelope (Prediction) that resembles the true staining (Groundtruth). The first row shows the x-y-plane, the bottom row the x-z plane of the 3D volume. (E) Example output for a classification task of benign and leukemic blood cells in blood smears from 200 individuals (C. Matek et al. 2019). We show two exemplary microscopy images (left) of two white blood cell classes, a monoblast and a neutrophil. The white blood cell type is predicted with a ResNet50. The confusion matrix (middle) shows that most of the 15 classes can be well predicted, in accordance to Matek et al. (2019).

### Pipeline Settings

After data preparation the user specifies tasks and parameters in the configuration file (see Fig. 1A). A maximum of eleven parameters have to be set. These are: The task (e.g., Semantic segmentation, instance segmentation, regression, or classification), the path to the project directory containing the train and test files, the pretrained weights for model initialization, the batchsize, the number of iterations over the dataset (e.g., epochs), the data augmentations, the loss function, the number of classes, if the images should be resized during import, if the uncertainty should be calculated after training, and if the model should be automatically evaluated after training. Within the pipeline one .json file serves as a config file.

After setting these parameters, the user executes the configuration file which starts the training with InstantDL on the desired task using the training data with the pre-trained weights and the chosen configurations.

### Transfer learning

Pre-trained weights can be used to initialize a training process, a practice called transfer learning. The choice of weights can have a huge influence on the performance of the algorithm: Transferring from natural images such as ImageNet to the medial domain seems not to be always beneficial (Ngiam et al. 2018) while using pre-trained weights from a medical dataset can improve performance (Hénaff et al. 2019). Pre-trained weights for 2D nuclei segmentation, 2D lung segmentation, 3D in-silico staining and for classification of white blood cells as well as ImageNet weights can be loaded.

### Data augmentation

Data augmentation is a method commonly used in machine learning to artificially increase the variance in the training dataset and thereby train the network to generalize better (Shorten and Khoshgoftaar 2019). We implemented spatial and color augmentations. The user can choose the desired augmentations, which are then randomly applied online, while importing the input images.

### Model training

InstantDL reads data from the corresponding folders and prepares for training and testing. This includes the following steps: i) Model initialization with pretrained weights, if selected. ii) Import of data and normalization, split of validation data from the data contained in the train data folder, shuffle of training data, batch creation and online data augmentation. iii) Training of the model for the given number of epochs using early stopping, which can be monitored live using tensorboard and automated saving of the best model. iv) Prediction of labels from the test dataset. v) For semantic segmentation, pixel-wise regression and classification uncertainty can be calculated after training.

### Model evaluation

The trained model is evaluated on the unseen test images and labels (i.e. the groundtruth). For semantic segmentation, instance segmentation and pixel-wise regression, the network predictions are saved as image stacks to ease evaluation of large datasets with limited CPU capabilities. This also allows an initial manual, qualitative evaluation and quality control. In a second step the predictions can be quantitatively evaluated. For that, accuracy, mean relative and absolute error, pixel-wise Pearson correlation coefficient and Jaccard index over all pixels of the test set are calculated. Boxplots to visualize quantitative model performance are generated. The standard quantitative evaluation output plots (i) the input images side-by-side to the corresponding labels and predictions and (ii) an error map between the labels and predictions to visualize training performance (see example evaluation Fig. 1B-D). For classification the predicted labels in the test set are compared to the true labels and multiple error scores (Jaccard index, mean absolute error, mean squared error, area under curve) are calculated. A confusion matrix and a receiver operating characteristic (ROC) curve are automatically visualized (Fig. 1E). All evaluation steps are implemented in the pipeline and can be set to be executed after testing. Additionally postprocessing (e.g. statistical analysis and visual assessment) is accessible in jupyter notebooks for customization, which are provided with InstantDL.

### Uncertainty quantification

Neural networks predictions can be unreliable when the input sample is outside of the training distribution, the image is corrupted or if the model fails. Uncertainty estimation can measure prediction robustness, adding a new level of insights and interpretability of results (Gal and Ghahramani 2016; Lakshminarayanan, Pritzel, and Blundell 2017). Bayesian inference can be approximated in deep Gaussian processes by Monte Carlo dropout (Gal and Ghahramani 2016). We have implemented Monte Carlo dropout for semantic segmentation, pixel-wise regression and classification in the pipeline. During the inference phase, 20 different models are created using Monte Carlo dropout and model uncertainty is calculated on the test set. For pixel-wise regression and semantic segmentation, the pipeline saves an uncertainty map. From this, pixel uncertainty can be plotted and saved to the project folder or average pixel uncertainty for an image can be calculated (Fig. 2A,B). For classification InstantDL adds the uncertainty score to the results file where a score close to zero indicates certain predictions, and score close to 1 indicates high uncertainty (Fig. 2C,D).

**Figure 2:**
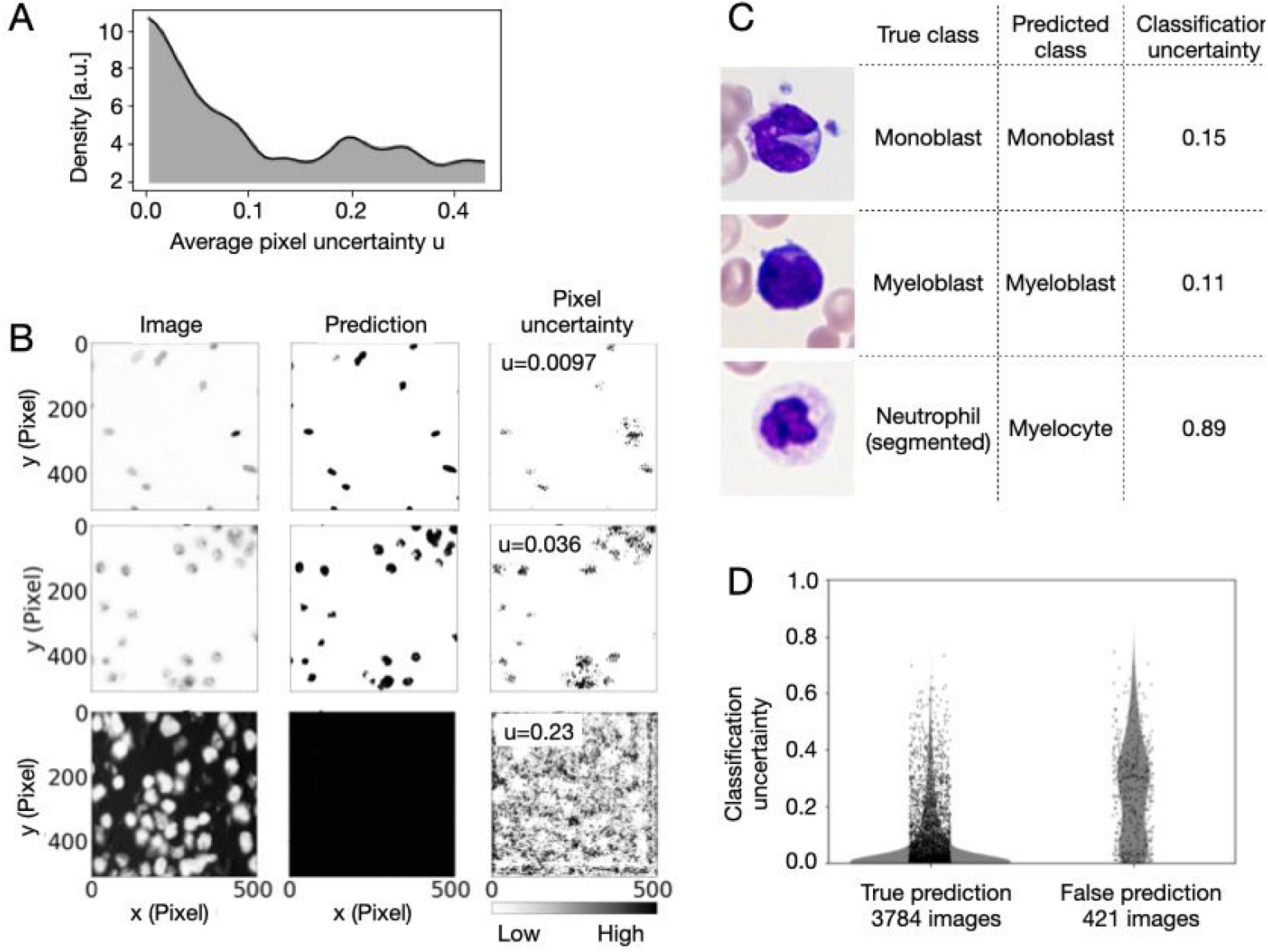
Uncertainty of semantic segmentation, pixel-wise regression and classification can be estimated with InstantDL. (A) The distribution of average pixel uncertainty u for 162 images of a semantic segmentation task (Caicedo, Goodman, et al. 2019). The distribution is approximately bi-modal. (B) Three exemplary semantic segmentations from the data visualized in (A) with InstantDLs pixel uncertainty estimation. Correct predictions correspond to a low average pixel uncertainty u (top and middle row), while a high average pixel uncertainty indicates failed segmentations (bottom row). Regions with ambiguous predictions are indicated by high pixel uncertainty (right column). (C) For the prediction of white blood cells classes (Christian Matek et al. 2019), classification uncertainty indicates incorrect predictions. (D) The distributions of classification uncertainty for correct and false predictions differ significantly (p-value < 0.001, Mann-Whitney rank test).

### Implementation

InstantDL is written in Python 3.6 and leverages Tensorflow 1.14.0 and Keras 2.1.5 (Chollet and Others 2015), which provide an excellent framework for our pipeline due to the modular, composable and user friendly design. The pipeline can be run locally ensuring data protection, on a cluster or with Google Colab (Bisong 2019), which has successfully been used for deep learning projects (“Deep Learning to Detect Skin Cancer Using Google Colab” 2019) making it usable for those with limited computational resources. We provide the pipeline as one package in a Docker image (“Docker Documentation” 2020) to simplify installation and application.

## 4 Results

To evaluate InstantDL broadly, we applied it to seven publically available datasets (four of which come from data science challenges) and compared its performance to published results. If no test set was provided, we took 20% of the data to create our own test set. This was done on the highest level of abstraction, for example on patient level or tissue-slide level whenever possible, otherwise the data was randomly split. We used the same evaluation metrics as published in the respective papers (Jaccard index, AUC, Pearson correlation) to compare our results appropriately.

For pre-processing, we transformed the images to .tiff files and classification labels to a .csv file to adapt them to the InstantDL requirements. Training was performed by saving the best model using early stopping. As data augmentation we used horizontal and vertical flipping. For pixel-wise regression we used mean squared error loss, for semantic segmentation we used binary cross entropy loss and for classification we used categorical cross entropy loss. For instance segmentation binary cross entropy was used as segmentation loss in combination with the localization and classification loss in the Mask-RCNN (Kaiming et al. 2017).

We evaluated the performance of semantic segmentation and instance segmentation on three datasets. In the first dataset we segmented nuclei in microscopy images contained in the Data Science Bowl 2018 (Caicedo, Goodman, et al. 2019) dataset. Using InstandDL instance segmentation we reached a median Jaccard index of 0.60 (25-75%ile: 0.61 to 0.58 estimated from bootstrapping), while using semantic segmentation we reached a median Jaccard index of 0.16 (25-75%ile: 0.15 to 0.17). The winner of the challenge reached a Jaccard index of 0.63 while the median participant reached 0.42 (solid and dotted line, Fig. 3A). The second task was the multi-organ nuclei segmentation challenge. Here, 30 microscopy images of various organs with hematoxylin and eosin staining are provided (Kumar et al. 2020, 2017). We reached a median Jaccard score of 0.57 (25-75%ile: 0.56 to 0.59) with InstantDL’s semantic segmentation and 0.29 (25-75%ile: 0.28 to 0.30) with instance segmentation. The winner of the challenge reached 0.69 and the median participant scored 0.63 (solid and dotted line, Fig. 3B). Thirdly, we benchmarked InstantDL on lung CT images from the Vessel-12 challenge (Rudyanto et al. 2014). Using instance segmentation we reached an area under the receiver operator curve (AUC) of 0.94 (25-75%ile: 0.94 to 0.94), and 0.90 (25-75%ile: 0.88 to 0.92) with instance segmentation. The winner of the challenge reached a score of 0.99 and the median participant 0.94 (solid and dotted line, Fig. 3C).

**Figure 3:**
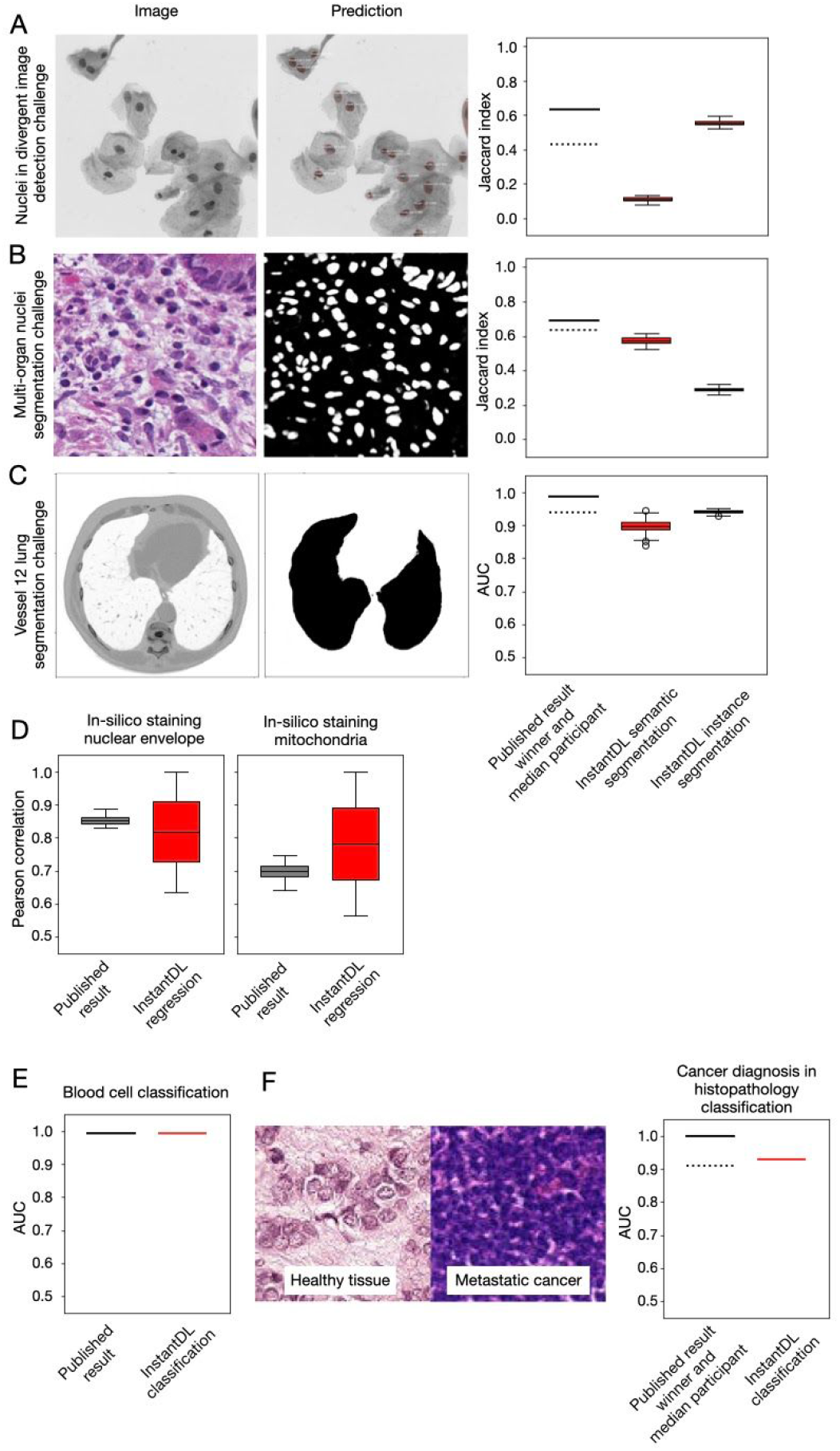
InstantDL achieves competitive performance on published datasets and computer vision challenges without hyperparameter tuning. (A) InstantDLs instance segmentation achieves competitive results on the nuclear detection challenge dataset (Caicedo, Goodman, et al. 2019), which contains a variety of experimental conditions, cell types, magnifications, and imaging modalities. We show one exemplary image from the dataset and the corresponding prediction using InstantDL’s instance segmentation. The winner of the challenge achieved a Jaccard index of 0.63 (solid line), while the median participant achieved 0.42 (dotted line). InstandDLs instance segmentation achieved a median Jaccard index of 0.60 without hyperparameter tuning. We estimate the Jaccard index distribution by bootstrapping, sampling 100 times half of the test set. Boxes indicate the median and the 25/75%ile of the distribution, whiskers indicate the 1.5 interquartile range. (B) For the challenge of segmenting nuclei in microscopy images of multiple organs with hematoxylin and eosin staining (Kumar et al. 2020, 2017), the winner achieved a Jaccard index of 0.69 (solid line) and the median participant 0.63 (dotted line). InstantDL using instance segmentation reached a Jaccard index of 0.29, and 0.57 using semantic segmentation. (C) Evaluation of instance segmentation of lung CT images from the Vessel-12 dataset (Rudyanto et al. 2014). The winner of the challenge reached an area under the receiver operating characteristic curve (AUC) of 0.99, while the median participant reached 0.94. InstantDL reached an AUC of 0.90 with semantic segmentation, and 0.94 with instance segmentation. (D) InstantDL’s pixel-wise regression performs similarly well as the published approach ((Ounkomol et al. 2018) for in-silico staining of bright-field images in three dimensions, but with a higher variability. We achieved a median pearson correlation of 0.85 for nuclear envelope staining and 0.78 for mitochondria staining. (E) For classification of leukemic blast cell images vs. benign white blood cell images (C. Matek et al. 2019; Shen et al. 2019), InstantDL achieved an AUC of 0.99, while Matek et al. report 0.99. (F) Classification of metastatic cancer in small image patches taken from larger digital pathology scans on histopathological images (“Histopathologic Cancer Detection” n.d.). InstantDL achieved an AUC of 0.93 while the winner of the challenge achieved an AUC of 1.0 and the median participant 0.91.

To evaluate InstantDL’s performance for pixel-wise regression, we predicted the 3D nuclear envelope and mitochondria staining from brightfield images (Ounkomol et al. 2018). For the nuclear envelope staining prediction, we achieved a median pixel-wise Pearson correlation to the real staining of 0.85 and 0.78 for the prediction of mitochondria staining (Fig. 3D), similar to the published result.

Classification performance was evaluated on two datasets. In Matek et al. images of single white blood cells from 100 leukemic and 100 non-leukemic individuals were classified into leukemic blast cell images vs. benign white blood cell subtype images (Christian Matek et al. 2019). We reached an AUC of 0.99 where Matek et al. reached 0.99 (Fig. 3E). The second task was to classify metastatic cancer image patches of digital pathology scans, taken from the Histopathologic Cancer Detection Kaggle challenge (“Histopathologic Cancer Detection” n.d.). We reached an AUC of 0.93, while the winner reached 1.00 (solid line in Fig. 3F) and the median participant scored 0.91 (dotted line, Fig. 3F).

## 5 Discussion

We present InstantDL, a deep learning pipeline for semantic segmentation, instance segmentation, pixel-wise regression and classification of biomedical images. InstantDL simplifies the access to the advantages of deep learning algorithms for biomedical researchers with limited computer science background. The only requirement is a solid understanding of the data (and how appropriately split it into training and test set), as well as of the task and loss function that should be optimized during training the model (see e.g. (Goodfellow, Bengio, and Courville 2016; LeCun, Bengio, and Hinton 2015)). The pipeline is designed for maximum automation to make training and testing as convenient and as easy as possible. However, some parameter settings depend on the dataset properties and therefore cannot be automated. After setting a maximum of 11 parameters, the pipeline can be run without further user interactions. We included state-of-the-art analysis metrics that are accessible out of the box. Moreover, we included uncertainty prediction to provide an additional level of interpretability of predictions. We tested the performance of InstatDL on a variety of publicly available data sets and achieved competitive results without any hyperparameter tuning.

Other deep learning frameworks such as OpenML (Vanschoren et al. 2013), Google Cloud AI (Yilmaz, Aydin, and Demirbas 2014), ImJoy (Ouyang et al. 2019) and ZeroCostDL4Mic (Von Chamier et al. 2020) provide similar tools for applying deep learning methods on user data (Table 2). However, they all require data upload to a cloud system. OpenML, e.g., offers an online ecosystem of datasets and machine learning models, but the dataset will be made publically available with upload. Google Cloud AI offers a framework for data analysis with predefined, not-modifiable models and a fee will be accrued. ImJoy and ZeroCostDL4Mic are browser based applications, requiring upload of the data to the web. Our pipeline can easily be installed and run locally on a computer or server, ensuring data privacy and security. Additionally it can be used on cloud solutions, such as Google Colab. As the code is publicly available it is convenient to tune and extend InstantDL.

**Table 1:** Overview on the image processing tasks implemented in InstantDL, required input, label, and output format.

|  | **Semantic segmentation** | **Instance segmentation** | **Pixel-wise regression** | **Classification** |
| --- | --- | --- | --- | --- |
| Input image | 2D & 3D | 2D | 2D & 3D | 2D |
| Labels | Binary images | Images with float pixel values | Images with float pixel values | Labels in a .csv file |
| Output | Binary images | Binary masks | Images with float pixel values | Labels in a .csv file |
| Architecture | U-Net | Mask RCNN | U-Net | ResNet50 |

**Table 2.** Comparison of InstantDL to other deep learning pipelines.

|  | **InstantDL** | **Open ML** | **Google Cloud AI** | **ImJoy** | **ZeroCostDL4 Mic** |
| --- | --- | --- | --- | --- | --- |
| **Host** | Local, on cluster, or Google Colab | Web based | Web based | Web based | Web based (Google Colab) |
| **Data privacy** | Yes, if run locally | No (shared with upload) | Limited (hosted in the cloud) | Limited (hosted in the cloud) | Limited (hosted in the cloud) |
| **Target audience** | Biomedical researchers | Researchers and developers | Enterprises | Researchers and developers | Biomedical researchers |
| **Developed for** | Biomedical images | All kinds of data | All kinds of data | All kinds of data | Biomedical images |
| **Customizability of Code** | Open source | Open source | Code not available | Possible through writing a plugin | Open source |
| **Cost** | Free | Free | Payment plan | Payment plan | Free |

Applied to microscopy images of single cells, deep learning-powered image processing can contribute to uncovering and understanding of a wide variety of molecular and cellular processes (Van Valen et al. 2016; Moen et al. 2019). With InstantDL, we hope to empower biomedical researchers to conduct reproducible image processing with a convenient and easy-to-use pipeline.

## 6 Availability of data and materials

InstantDL is published on GitHub: https://github.com/marrlab/InstantDL. There, we provide code, extensive documentation, and instructive examples of how to run the pipeline.

For reproducing our results, all data used for illustration and benchmarking is available for download at https://hmgubox2.helmholtz-muenchen.de/index.php/s/YXRD4a7qHnCa9x5

## 8 Authors’ contributions

Dominik Waibel implemented the pipeline and conducted experiments. He wrote the manuscript and created the figures with Carsten Marr and Sayedali Shetab Boushehri. Sayedali Shetab Boushehri revised and refactored the code, added docker installation and tested the pipeline. Carsten Marr supervised the study.

## Acknowledgements and Funding

We thank Niklas Köhler and Nikos Chlis (Munich) for contributing to the initial concept for InstantDL. We thank Daniel Schirmacher (Zürich), Lea Schuh, Johanna Winter, Moritz Thomas, Matthias Hehr, Benjamin Schubert, Niklas Kiermeyer, and Benedikt Mairhörmann (all Munich) for valuable feedback on the manuscript and using InstantDL. This project has received funding from the European Union’s Horizon 2020 research and innovation programme under grant agreement No 862811 (RSENSE). Carsten Marr was supported by the BMBF, grant 01ZX1710A-F (Micmode-I2T). Sayedali Shetab Boushehri is a member of the Munich School for Data Science (MUDS).

